# Sharks, Lies, and Videotape: A content analysis of 32 years of Shark Week documentaries

**DOI:** 10.1101/2021.08.18.456878

**Authors:** Lisa B. Whitenack, Brady L. Mickley, Julia Saltzman, Stephen M. Kajiura, Catherine C. Macdonald, David S. Shiffman

## Abstract

Despite evidence of their importance to marine ecosystems, at least 25% of all chondrichthyan species are estimated or assessed as threatened with extinction. In addition to the logistical difficulties of effectively conserving wide-ranging marine species, shark conservation is believed to have been hindered in the past by public perceptions of sharks as dangerous to humans. Shark Week is a high-profile, international programming event that has potentially enormous influence on public perceptions of sharks, shark research, shark researchers, and shark conservation. However, Shark Week has received regular criticism for poor factual accuracy, fearmongering, bias, and inaccurate representations of science and scientists. This research analyzes the content and titles of Shark Week episodes across its entire 32 years of programming to determine if there are trends in species covered, research techniques featured, expert identity, conservation messaging, type of programming, and portrayal of sharks. We analyzed titles from 272 episodes (100%) of Shark Week programming and the content of all available (201; 73.9%) episodes. Our data demonstrate that the majority of episodes are not focused on shark bites, although such shows are common and many Shark Week programs frame sharks around fear, risk, and adrenaline. While anecdotal descriptions of disproportionate attention to particular charismatic species (e.g. great whites, bull sharks, and tiger sharks) are accurate and supported by data, 79 shark species have been featured briefly at least once. Shark Week’s depictions of research and of scientists are biased towards a small set of (typically visual and expensive) research methodologies and (mostly white, mostly male) scientists, including presentation of many white male non-scientists as experts. While sharks are more often portrayed negatively than positively, limited conservation messaging does appear in 53% of episodes analyzed. Results suggest that as a whole, while Shark Week is likely contributing to the collective perception of sharks as monsters, even relatively small alterations to programming decisions could substantially improve the presentation of sharks and shark science and conservation issues.

## Introduction

Shark species can make important contributions to the resilience, structure, and function of marine ecosystems (1, 2). The loss of shark populations has the potential to influence prey abundance, diversity, behavior, and genetics, although the nature and extent of that influence varies based on ecological context (reviewed in Ferretti et al. 2010). At least 24 percent of all chondrichthyan species (sharks, rays, skates, and chimeras) are assessed as or estimated to be Threatened with extinction (3–5), primarily due to overfishing (6, 7). Conservation of shark populations is challenging given life history characteristics including slow growth, late age at maturity, and relatively low fecundity (8). Management is further complicated by the reality that shark fisheries globally are often under-managed, enforcement resources are typically limited, many species are highly mobile through multiple management jurisdictions, and sharks can represent an important food source, especially in subsistence fishing communities (3).

In addition to the biological, ecological, and practical difficulties of effectively conserving sharks, shark conservation is believed to have been hindered in the past by public perceptions of sharks as dangerous to humans (9), including portrayals suggesting that they are evil or vicious (10).

One of the primary ways that the public (defined here as people who are not shark scientists, marine scientists, or aquarists) obtains their knowledge about sharks is through the media, including news stories, social media, and television programs (11, 12). Although negative images of sharks predate the modern era, with sharks featured as villains in many famous works of art and literature (e.g. Copley’s *Watson and the Shark* (1778) or Hemingway’s *The Old Man and The Sea* (1952)), the movie *Jaws* (1975) marked a shift towards modern presentations of sharks with a visceral visual immediacy. These negative portrayals have been reinforced over decades by media reporting focused overwhelmingly on shark bite incidents and by “attack” focused movies and television programs (10,13–15). This media landscape contributes to a collective public conceptualization of sharks as bad, dangerous man-eaters (16, 17).

Public perceptions of sharks may affect shark conservation efforts directly or indirectly, by altering public attitudes or by shaping public knowledge and support for particular policies (9, 18). In addition to a primary focus on covering shark bites (14), reporting on sharks in newspapers focuses attention on particular threats to sharks and disproportionately discusses a relatively small number of charismatic species, regardless of conservation status--potentially leading to failures to direct attention to the most serious conservation challenges or most threatened species (19). In a participatory democracy, conservation policy change often requires public support, and fame is likely to have both costs and benefits in terms of conservation attention and support (18, 20).

Shark Week, an annual event from Discovery Communications, has played a large role in shaping public perceptions of sharks since 1988. Shark Week is the highest-profile coverage of marine biology or ocean conservation on American television, and represents the greatest temporary increase over baseline in Americans paying attention to any ocean science topic (21, 22). In 2020, over 21 million viewers tuned in and 37% of those viewers did not watch the Discovery Channel in the month prior to Shark Week (23). Social media mentions occur in the hundreds of thousands each year during Shark Week, as do notable spikes in Wikipedia searches about sharks (21). Therefore, this long-running programming event has potentially enormous influence on public perceptions of sharks, shark research, shark researchers, and shark conservation; it may be the only time that many people think about these topics at all during a typical year.

Shark Week has received regular criticism for poor factual accuracy, fearmongering, bias, and inaccurate representations of science and scientists (e.g. (24–27). Multiple scientific experts have reported that their words were selectively edited to make it appear that they were responding to questions they were not asked, or that they found the way they were presented in Shark Week programs profoundly professionally embarrassing (28, 29). In 2010, in response to criticism, Discovery Communications agreed to reduce entertainment programming during Shark Week and present more scientifically-oriented episodes, though many of these more scientific programs continued to depict sharks negatively (30).

Fictional programming from Shark Week has also generated scientific criticism, with shows like *Megalodon: The Monster Shark Lives* (2013), *Megalodon: The New Evidence* (2014), and *Shark of Darkness: Wrath of Submarine* (2014), presenting fictional storylines featuring CGI and actors pretending to be scientists and government officials, without clear communication to the viewer that these programs are not factual (e.g. (31, 32). According to a post-show social media poll following *Megalodon: The Monster Shark Lives,* 79% of respondents reported believing that *Otodous megalodon* was still alive (31), despite the fact that all scientific evidence suggests that *O. megalodon* went extinct over 2 million years ago (33, 34). These types of programs not only present inaccurate information in ways that may reduce trust in science and scientists (e.g. (35), but through their depiction of scientific and government collusion to hide the truth, join broader sociological trends towards belief in conspiracy theories, which can undermine confidence in previously undisputed facts and limit the potential for social consensus-building around a wide range of important issues (36).

Defenses of Shark Week have typically fallen into two primary categories: firstly, that the most problematic elements of Shark Week are also essentially harmless (e.g., that no-one expects Shark Week to offer accurate information or believes what they see there (Shiffman, pers. obsv.)). Not-yet-published data suggests that this is not the case, and that the public generally perceives Shark Week as an accurate source of information about sharks (Wester et al. pers. comm.). It is well established that reality television and info- or edu-tainment programming can have real-world effects on public understanding of issues. Evidence suggests that exposure to pseudoscientific television programming predicts pseudoscientific beliefs (e.g. (37, 38), and that regular exposure to reality TV featuring the paranormal predicts endorsement of paranormal beliefs (39). Sixty-eight percent of pregnant women reported watching reality television shows about pregnancy and birth, and 72% of those women who were pregnant for the first time said that reality television could help them understand what it was like to give birth, although these shows disproportionately focus on medically- and technologically-assisted birth, present birth as dangerous, and frame women’s bodies as incapable (40, 41). Reality TV filming locations have been reported to have substantial effects on tourist destination choice and demand for particular locations (42). Television viewing shapes adolescent’s beliefs about the risks of alcohol consumption and their intention to drink (43). The power of media has become especially apparent this past year during the COVID-19 pandemic, as 2020 was marked by the propagation of medical disinformation and conspiracy theories across social media, with an estimated 30% of the U.K. and U.S. populations subscribing to COVID scientific conspiracy narratives (44). These and many other similar examples refute arguments that Shark Week programming can have no meaningful real-world impact on sharks, shark science, or shark conservation.

Secondly, defenders of current programming have argued that alternatives do not exist (e.g., that it is impossible to include more diverse representations of shark scientists because available shark scientists are all white men (27) or that audience demand will not allow for more conservation or science-focused programs (45)). These arguments are undermined by the existence of organizations like Minorities in Shark Science (MISS;(46)) which to date has over 300 members that identify as women of color (Graham, pers. comm.), and the popularity of factual, documentary television nature programming like that found on the BBC (as opposed to the more “hybrid” programming combining factual and non-fiction or fictional genres (47) often seen on Shark Week). While viewership for the Blue Planet II series was notably lower in the U.S. than in the U.K., each episode had approximately 3 million viewers (48).

Shark Week represents a significant opportunity for public engagement with shark science and conservation for a large and enthusiastic audience. Between 2014 and 2017, the Twitter discourse regarding sharks shifted from negative or slightly negative to slightly positive or positive during Shark Week air dates compared to non-Shark Week parts of the year (49).

Documentary programming can shape public opinion about conservation-relevant issues, particularly in concert with other reinforcing information or events, sometimes resulting in pro-environmental policy and management changes (50). With adjustments to some choices about programming, Discovery Communications could substantially improve their messaging about science, sharks, and shark conservation and increase representation of diverse scientists on television.

The goal of this research was to quantify some of the reported trends in Shark Week programming, analyze the content and titles of Shark Week episodes to determine if there are trends in species covered, research techniques featured, expert identity, conservation messaging, type of programming, and portrayal of sharks. This quantitative analysis of the key features of Shark Week’s 30+ year run will provide better data from which to discuss current practices and recommend improvements.

## Methods

### Title Analysis

Titles from all Shark Week episodes from 1988 to 2020 were collected from two online sources: thetvdb.com (2021) and the Washington Post (1994) (N=272). Each title was subjectively analyzed to determine whether it contained words evoking negative connotations or if the title could be taken as negative within context. Individual title words were searched within the list of Affective Norms for English Words (ANEW) (51). Title words that scored below the mean for valence (ranging from pleasant to unpleasant) or above the mean for arousal (ranging from calm to excited) were considered as eliciting a negative effect. These titles often included words associated with violence (wrath, fury), fear (terror, fear, scream), death (deadly, death, killer), danger (danger, dangerous), or attacks (shark attack, shark bite).

In addition to analyzing individual words within titles, whole titles were analyzed in context. This allowed some titles that contained no negative words from the ANEW list to be categorized as negative based upon the construction of the title, such as “Rogue Shark”, “Jaws Strikes Back”, or “Lair of the Mega Shark”. Titles that contained the word “Jaws” were treated as negative since the word was deliberately chosen based on its association with the eponymous movie.

Episodes from the “Air Jaws” series were not treated as negative unless other parts of the title had a negative connotation, as this series already has a reputation for not being negative. The list of titles was independently analyzed by a public relations professional in the same manner to confirm our subjective analysis.

The total number of titles, and number of negative titles was counted for each year and the proportion of negative titles was calculated. Each title was stripped of punctuation and decomposed into individual words. Singular and plural words were grouped together (eg. Shark, Sharks) and root words with various suffixes were also grouped together (eg. Kill, Killer, Killing).

The frequency of occurrence of words and word groups was sorted from most to least frequent for all years combined. Lines of best fit were calculated in Excel®.

### Episode Analysis

For the purpose of this study, obtainable episodes (N=201, 73.9% of all aired Shark Week episodes; Supplement 1) of Discovery Shark Week® from 1988-2020 were watched by four trained coders (LBW, BLM, JS, and DSS). Episodes which were analyzed were obtained from a variety of online sources (i.e., Hulu®, Amazon Prime®, Discovery+®, YouTube®, Vimeo®), library loans, and private holdings. Episodes from earlier years were watched on VHS tape. Those which were not coded were not obtainable via any of the aforementioned platforms, despite extensive online searches between 2019 and 2021.

A code book was developed by all coauthors, which is described in detail in Supplement 2. The coding process included the following areas of interest: A) documentary title, B) documentary year, C) production company, D) locations of filming, E) species of chondrichthyans (sharks, rays, skates, chimeras) featured, F) featured experts/hosts of the show, G) general type of documentary, H) research methods featured, I) purpose/goal of documentary, J) accomplishment of goal/purpose, K) mention of shark conservation, L) mention of shark finning, M) mention of shark meat, N) mention of ways to help sharks, O) negative portrayal of sharks, P) positive portrayal of sharks, Q) portrayal of sharks other than negative or positive, R) mention of specific misconceptions (e.g., “bull sharks are the only sharks that can enter freshwater” and “sharks can smell a drop of blood from a mile away”), S) anything else about a given episode that coders found noteworthy including space for coders to note variables that they were unsure about. Most variables were binary (presence/absence), which minimized the chance of coding error or coding agreement problems resulting from interpretation.

Prior to data collection, all coders completed analysis of the same three episodes to check for intercoder reliability and assess whether the coding tool was working as intended. There were no differences in quantitative coding or results. Coders filled out the same structured coding form in Google Forms® for each episode, filling in applicable information throughout the duration of the show. Each episode was coded by one coder. Only one coder (LBW) appeared in an episode, which was assigned to a different coder. A single coder (DSS) acted as arbitrator to minimize error resulting from coder uncertainty and to make decisions on any variables that were coded unclearly.

### Hosts/Expert Analysis

Not-yet-published data from a recent survey done by Wester et al. indicate that Shark Week is perceived by the public as a reputable source of information on sharks and conservation (Wester, pers. comm.). To better understand the sources of expert perspectives offered by Shark Week, we assessed the academic research productivity of those identified onscreen as “experts”, “scientists”, and “researchers” as a rough proxy for recognized scientific activity. We used Google Scholar to determine the number of peer-reviewed publications (as of July 9, 2021) authored by each Shark Week expert/scientist. Because scientific journal publications are not the only evidence of scientific expertise, we also counted authorship of peer-reviewed book chapters and edited books, peer-reviewed abstracts from conferences, and government reports. Popular press books and magazine articles were not included. We sorted expert scientific productivity into bins based on number of publications (0, 1-5, 6-15, 16-25, 26-50, 50+ publications). Fictional experts were removed from the analysis (i.e. fictional scientist “Colin Drake” from the megalodon episodes). In order to assess the gender and racial diversity of non-fictional hosts/experts, we conducted Google® searches for biographies, news articles, or social media profiles of each host/expert.

We successfully located sources for all 204 non-fictional hosts/experts. The pronouns used in these sources were used to determine their gender, with the caveat that this allows us to speak only to broad trends in gender representation on Shark Week, not to the individual gender identities of particular experts. We recognize that people may be misgendered in sources that they did not write themselves.

We assessed host/expert race based on how they would likely be perceived by U.S. audiences, with the exception of the small number of hosts/experts who mention their race in public biographies or platforms (i.e., on Twitter #BlackinMarineScience or #LatinxinSTEMWithout).

Because names are a poor predictor of racial identity, we did not consider names in this analysis. Without drawing any conclusions about individual identities, this allowed us to develop an approximate estimate of the proportion of hosts/experts who can be assessed as white or white-passing.

### Species Analysis

After all episodes in the study were coded, we collected the following additional information on each shark species appearing in the episode: A) maximum recorded size (Castro, 2011), B) IUCN Red List Status (iucnredlist.org), C) taxonomic order. All species appearing on screen and also mentioned by name in narration were recorded, regardless of total screen time and regardless of whether that species was discussed meaningfully during the episode, thus results over-represent the coverage of less commonly featured species (e.g. a blacknose shark that appeared for ten seconds in an episode largely about tiger sharks would yield the same results as an episode that featured both blacknose and tiger sharks equally).

### Assessment of Scientists Attitudes toward Shark Week

A survey of expert shark researcher’s perspectives on shark conservation policy (51) included questions concerning perspectives on the role of media coverage in shaping public understanding of sharks. These results were collected with the approval of the University of Miami Human Subjects Research Office Institutional Review Board protocol 20130730, but have not previously been published. In this survey, experts are defined as members of professional scientific societies focusing on sharks and their relatives, including the American Elasmobranch Society, the Oceania Chondrichthyan Society, and the European Elasmobranch Society.

Questions whose answers we report here are “How significant do you feel that public attitudes towards sharks are in terms of shaping public policy?” and “In your opinion, does the mainstream media accurately portray shark science and conservation issues?”

## Results & Discussion

### Title Analysis & Programming Trends

Between 1988 and 2020, there were 272 unique Shark Week program titles (Supplement 1). For the first 15 years the number of original programs each year did not exceed seven. The number of programs began to increase dramatically after 2010, with the greatest number of unique programs (24) produced in 2018 and 2020 (Fig 1). Over half (51.8%) of all Shark Week programming was produced in the past eight years (2013-2020). The increase in the number of titles each year is best fit with an exponential function (y = 2.290e^0.0622x^ where x = year 1-33, r^2^ = 0.882). Overall, 59 titles (21.7%) used words with negative connotations, based on the ANEW. Although the number of titles produced increased exponentially, the number of titles with negative words each year ranged between zero and five and increased less dramatically (y =1.204e^0.039x^ where x = year 1-33, r^2^ = 0.449). When titles were analyzed within context, 42.6% of all titles were categorized as negative. The number of negative titles increased proportionally with the total number of titles as an exponential function (y = 0.938e^0.074x^ where x = year 1-33, r^2^ = 0.850).

**Fig 1.**
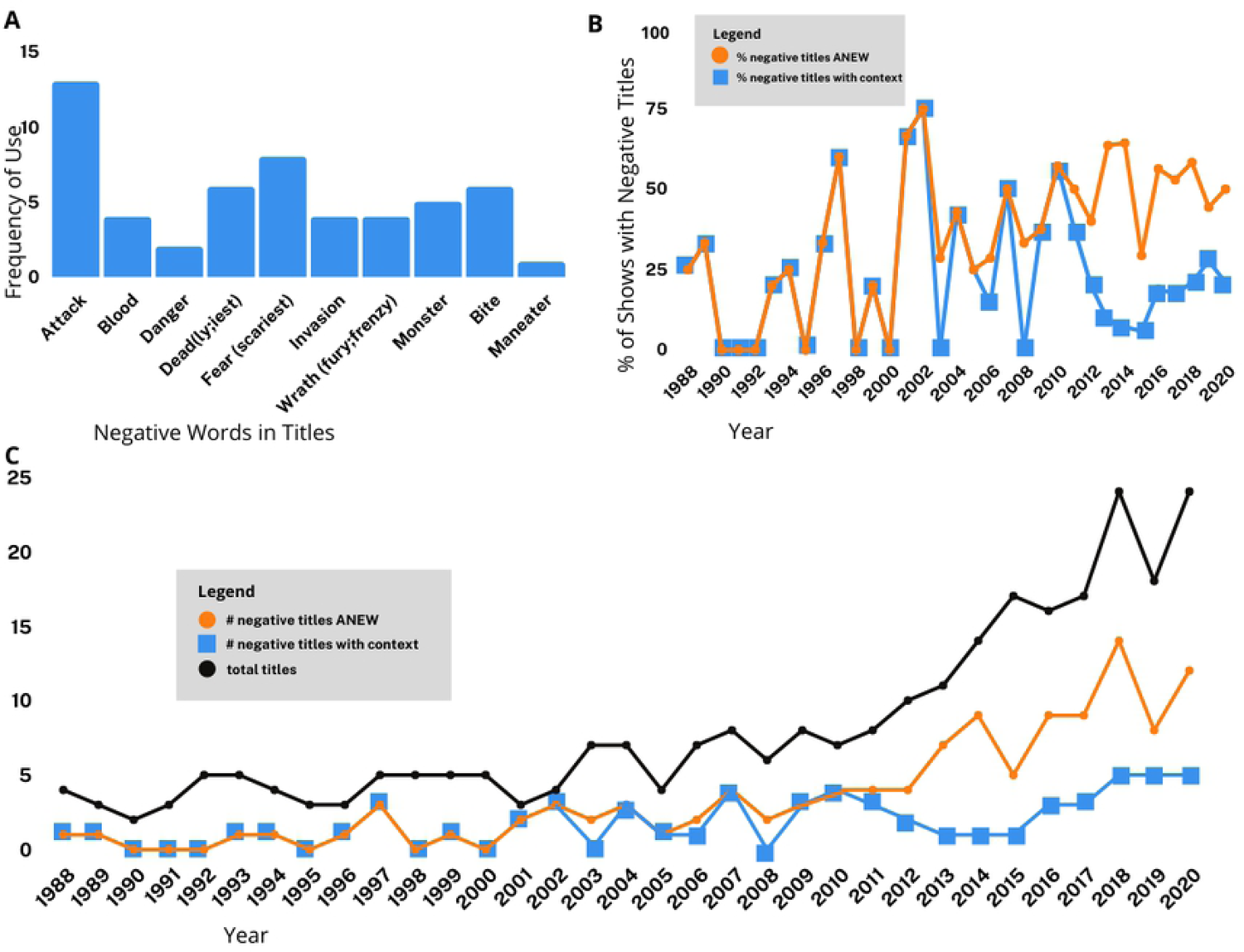
Trends in Shark Week episode titles. (A) Frequency of negative words used in episode titles; (B) Percent of episodes with negative titles per year. Orange circles indicate episode titles coded as negative based solely on the ANEW, blue squares indicate episode titles coded as negative by authors based on context; (C) Number of episodes with negative titles per year. Black circles indicate total number of episodes aired each year, orange circles indicate episode titles coded as negative based solely on the ANEW, blue squares indicate episode titles coded as negative by authors based on context.

The proportion of titles that were assessed as having negative connotations ranged from 0-75% in any given year (Fig 1). During the first five years of programming (1988-1992) only two titles were considered negative. In total, six non-consecutive years, (1990, 1991, 1992, 1995, 1998, 2000), representing 18.2% of the time examined, had no negative titles. The last year in which there were no negative titles was 2000. In contrast, at least half of the program titles each year were considered negative for twelve years, which represents 36.4% of the time examined. The years in which at least half of the program titles were negative were:1997, 2001, 2002, 2007, 2010, 2011, 2013, 2014, 2016, 2016, 2018, and 2020. The greatest number of negative titles was found in 2018 (14) and 2020 (12). These two years both produced the greatest number of programs (24 each year) and combined they represent 17.6% of all Shark Week programming since inception.

Title length ranged from one to eight words, with four word titles occurring most frequently. One hundred seventy-three titles (64.6%) included the root word “shark”. While 99 of the titles (36.4%) did not include the root word “shark”, many of these titles referred to a specific species, such as “Great White Encounters” or, “Search for the Golden Hammerhead”. Sixteen titles (5.9%) were merely descriptive and used the format, “Sharks of xxx”, where the xxx denotes a location.

The 272 program titles were composed of 1047 total words. There were 354 unique words, including root words plus their derivatives. The words “shark” and “sharks” occurred the most frequently (161 occurrences) and accounted for 15.4% of all title words. This was followed by the prepositions “of” (6.5%) and “the” (5.6%), then “jaws” (3.2%), “great” (2.8%), “white(s)” (2.6%), and “attack(s)” (1.4%). “Attack(s)” was the most frequently occurring word that has a negative connotation based on the ANEW list, and “jaws” was the most frequently occurring word that has a negative connotation in context (Fig 2).

**Fig 2.**
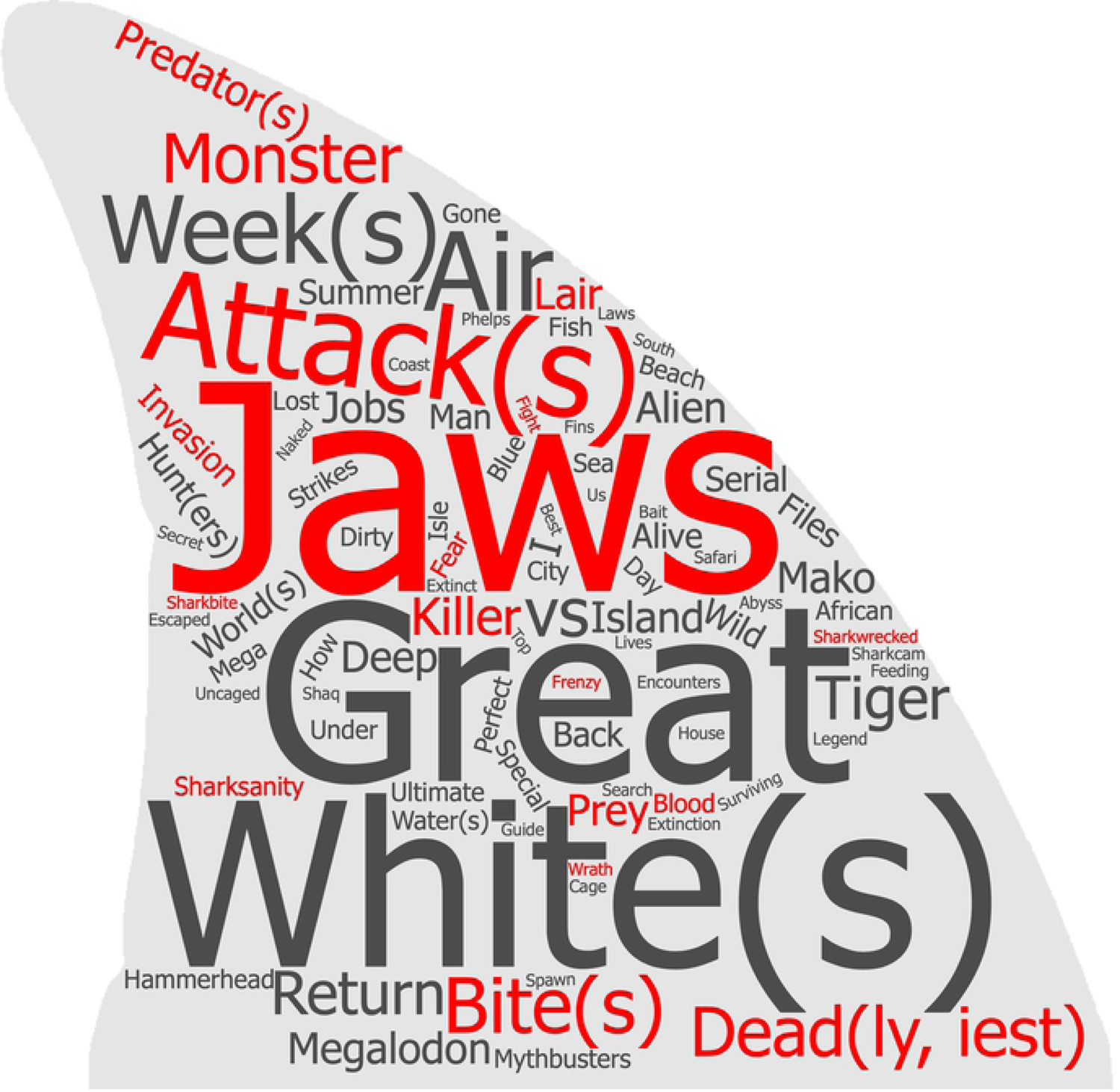
Frequency of occurrence of title words. The word “Shark(s)” occurred 4.7 times more frequently than the next most common word, “Jaws”, and was omitted from the word cloud. Prepositions were also omitted, along with words that occurred only once. Words with a negative connotation are depicted in red and font size reflects frequency of occurrence.

The number of negative titles for Shark Week shows increased at a lower rate (for words from the ANEW list) or at a similar rate (for titles taken in context) to the total number of titles. The number of negative titles based on the ANEW list did not exceed five per year, despite a dramatic increase in the total number of programs produced. For the first 15 years, all programs with negative titles were derived from words within the ANEW list. However, more recently, the number of negative titles in context has increased at a greater rate than the number of titles with negative words from the ANEW list. This suggests that titles are being constructed to avoid negative words, but are still depicting sharks in a sensationalized, potentially negative light when taken in context.

Some words that have a negative connotation in isolation can be rendered neutral in context. For example, an existing franchise entitled, “Naked and Afraid” produced episodes for Shark Week in 2018 and 2020 with the title, “Naked and Afraid of Sharks”. In this case the franchise title already included the word “afraid” so these titles were not included in the list of negative titles. However, taken in isolation, the title “Naked and Afraid of Sharks” would be considered to be a negative title. Similarly, the word “monster” is recognized as negative, but a program entitled “Monster Garage: Shark Boat” used the word “monster” to describe a garage, not sharks. In this case the title was not classified as negative. The word, “monster” can also refer to something that is very large, so programs like, “Monster Mako”, can refer to a particularly large mako shark. However, the word “monster” in this and similar titles was likely chosen to elicit fear and titles which used words like “monster” to describe sharks themselves were classified as negative despite some ambiguity around their meaning.

The word “Jaws’’ is unique in the context of Shark Week titles. Nearly all vertebrates possess jaws, so the word is not considered negative in itself. However, the 1975 movie “Jaws” caused people to associate the word with a killer shark. The word thus evokes a primal fear of being attacked, bitten, or eaten. Titles that include the word “Jaws” take advantage of this association by indirectly suggesting that the subject is dangerous or fear-inducing, without having to use words that are explicitly negative. Therefore, titles that included the word “jaws”, other than “Air Jaws”, were classified as negative within context.

There has been a recent trend to amalgamate “shark” with root words that have a negative connotation. This creates chimeras such as Sharkzilla (2012), Sharkpocalypse (2013), Sharkageddon (2014), Sharksanity (2014, 2015, 2016), and Sharkwrecked (2018, 2019). None of these fabricated words would appear in the ANEW list so they were analyzed within context. While the root words “apocalypse”, “armageddon”, and “shipwreck” are not included in the ANEW list, the root word “insane” is listed and does have a negative connotation (52).

### Episode analysis

#### General content

201 episodes were watched, coded, and scored, though not every variable was present in every episode. A plurality of episodes were broadly categorized as being about “Research” (37%) or “Natural History” (16%) (Fig 3). Our definition of research was very broad and essentially included any attempt to obtain the answer to any question about shark biology or behavior via observation or experimentation. Many of the episodes categorized as research include atypical or unscientific methods, attempts to answer questions long considered resolved in the peer-reviewed scientific literature, or experimental design that would likely not be considered scientifically valid if presented in an academic journal or conference. Episodes that focused on reenactments of sharks biting people (“Shark bites”) and episodes with no purpose beyond people diving with sharks (“”Diving with sharks”) each represented about 15% of all episodes.

**Fig 3.**
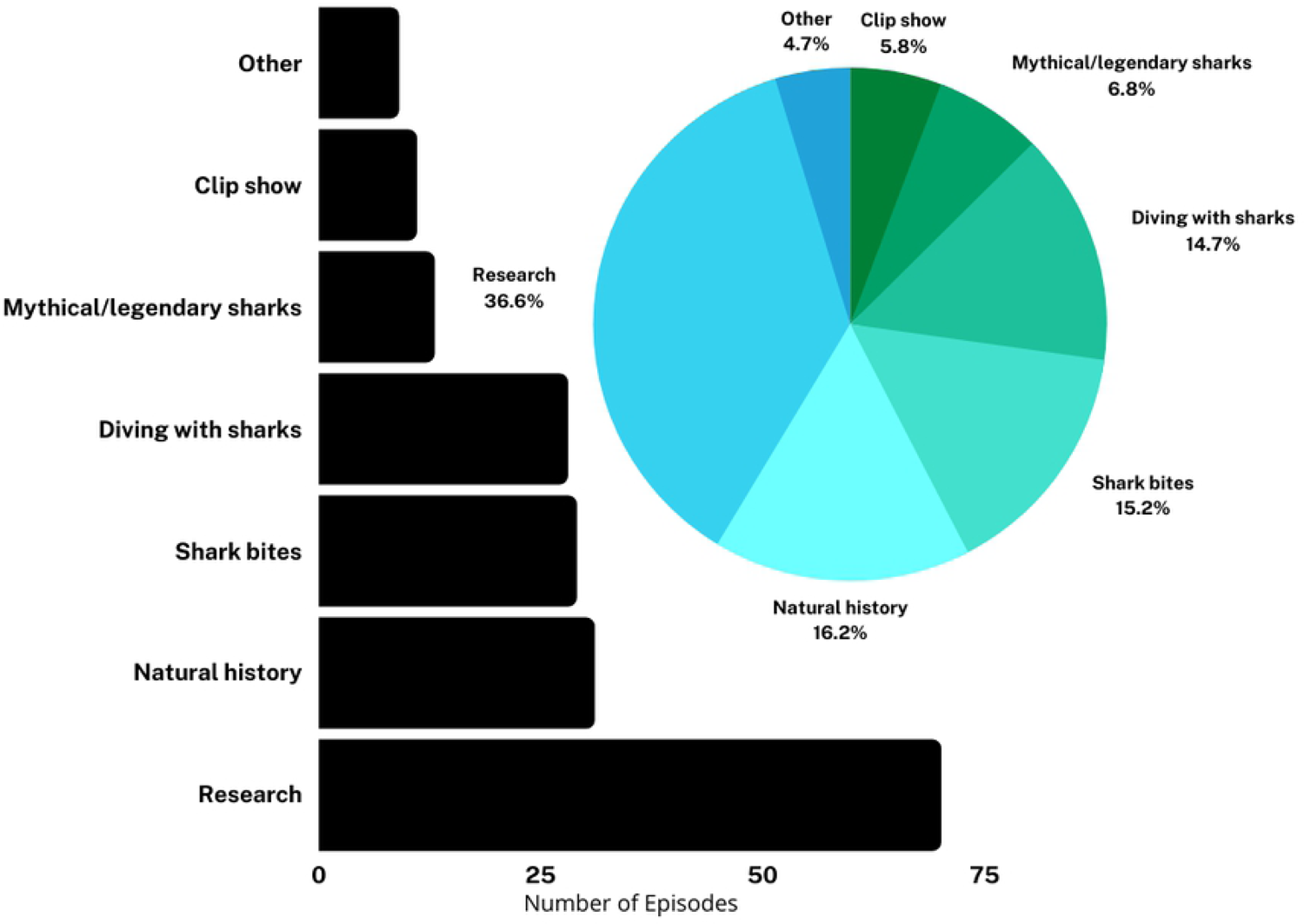
Number and percent of episodes by documentary class.

Episodes about mythical/legendary sharks represented about 7% of all episodes. When analyzed by year, we found no trends in programming; episodes have not become more or less focused on science or shark bites over time (Fig 4). However, we note that “research” themed programming was 25% or less in several years, including several consecutive years leading up to the fictional megalodon episodes (2009-2012) and the most recent year analyzed in this study (2020).

**Fig 4.**
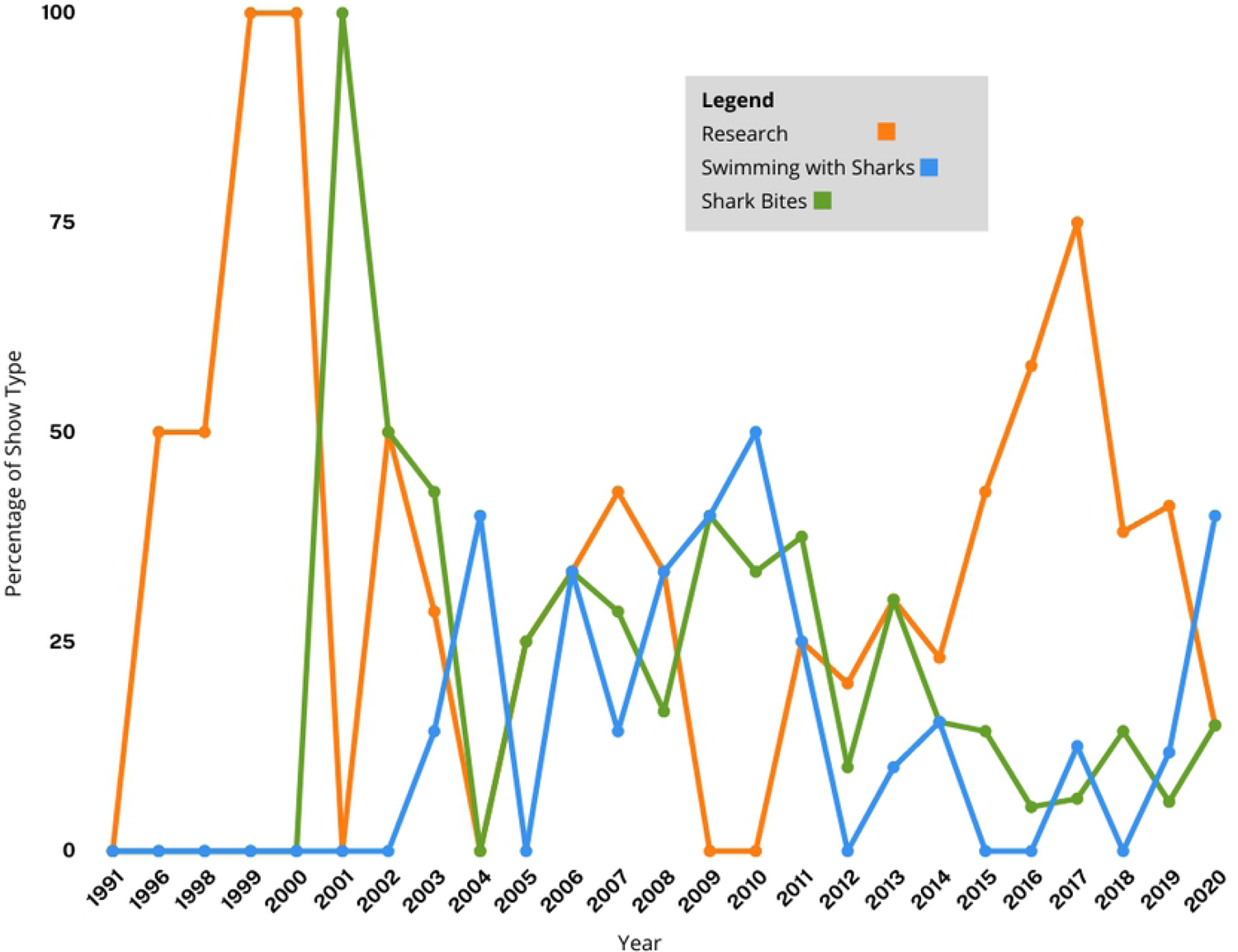
Percent of episodes categorized by “Research” (orange line), “Shark Bites” (green line), or “Swimming with Sharks” (blue line).

A focus on shark bites, shark-related danger, and mythical, legendary, or fictional monster sharks reflects, at least in part, the use of violence or fear as a marketing tool. Violent programming is a market differentiator known to attract advertiser-desired demographics, particularly 18-34 year old males (53). Hamilton (53) describes this tendency as an economically rational and self-interested act by networks, despite creating negative externalities for society that are not borne by the producers or programmers who are making decisions about content. It is also possible that the creation of frustrated and vocal constituencies opposed to inaccurate and fear-mongering programming, including opposition from scientists and conservationists, is part of an overall marketing strategy in which critique drives further public attention and viewership to even highly problematic content. Controversy and social media discussion are strongly positively correlated with sales performance, although strong and consistent negative word of mouth feedback may harm perceptions of a brand (54). This argument is supported in the context of Shark Week in particular by O’Donnell’s observation that the year that aired the fake megalodon documentary (2013) was also the year that generated the greatest volume of Twitter discussion about Shark Week (49).

### Research Methods

18.3% of Shark Week episodes featured no real research methods of any kind, which is the most common “research” category (Fig 5). Among episodes that did include research methods, methods featured tended to be simple, declarative and visual, such as satellite tagging large charismatic animals. The most common methods were satellite telemetry tagging, acoustic telemetry tagging, or use of high-tech camera equipment including drones, ROVs, BRUVs, or ultra-high-speed cameras; when added together these techniques were featured in approximately 40% of episodes. Some of this high-tech camera equipment is non-standard in published scientific research and was used in these episodes to obtain high-quality imagery for television, rather than scientific research purposes. While we note that examples of published studies using high-tech camera equipment does exist (as reviewed in 55), the definition of “research method” used here was inclusive of activities that would not meet the scientific threshold for “research”.

**Fig 5.**
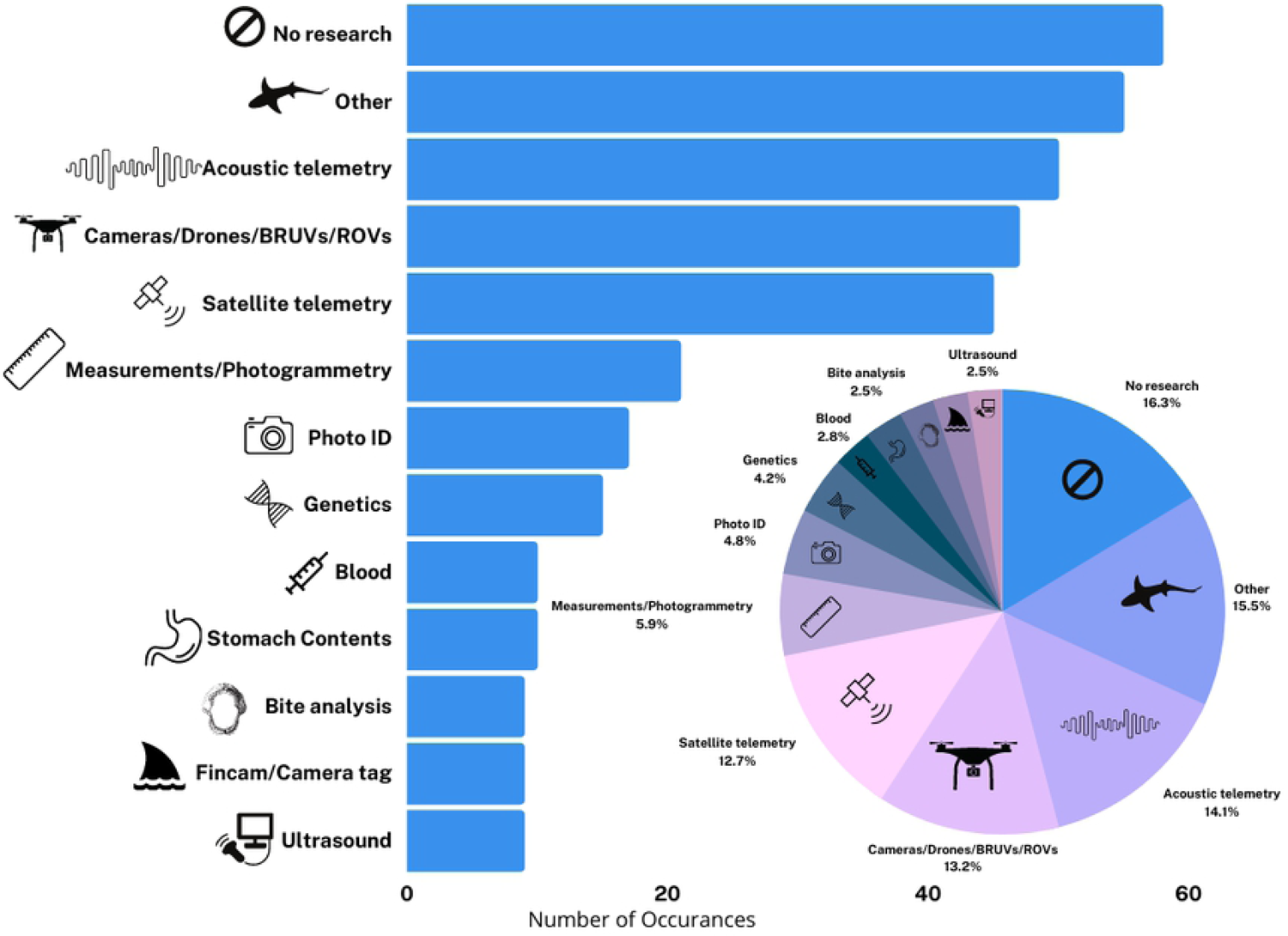
Number and percentage of occurrences of particular research methods in Shark Week episodes.

Research with high-tech equipment such as ROVs and satellite telemetry tagging is also expensive, which translates into featuring well-funded and prominent (often white and male) researchers. These research methodologies typically require more grant, donor, or personal funding, which is easier to access for scientists with some degree of seniority and influence and institutional support. Other methodologies producers consider less visually appealing may be more likely to be performed by scientists working with limited resources, including early career researchers, scientists at less wealthy institutions, or those from less wealthy countries, making them less likely to be featured in Shark Week episodes. Compensation for Shark Week appearances may also exacerbate these differences; scientists requesting industry-minimum pay for their appearances can be passed over in favor of those who have more financial resources and therefore may not need the additional funds (Jewell, pers.comm.). Appearing on Shark Week programming can have positive benefits for researchers, including increased visibility at home institutions and in the media, increased professional opportunities, and additional research funds or resources (56). The research methods featured in Shark Week are also notably distinct from the most common methods used in scientific investigations (57), which are dominated by age and growth, life history, and reproductive biology work (although see (55) for a review on the use of drones in shark research) While perhaps not as camera friendly, this kind of work is vital to generate data for the sustainable management of shark species, and the fact that commonly conducted, management-relevant science is rarely featured could impact public understanding of the purpose, function, and social relevance of marine science and the scientific process.

In addition to focusing on a very limited range of existing research techniques, television programming often presents science and scientific discovery as reporting unquestionably true facts rather than as generating knowledge through human-led iterative processes (58). A significant disconnect has been found between scientists (who generally described science on television as failing to reflect the practices and methods of science), and those working to produce science programs for television, who believed reflecting uncertainty and methodological processes in science television programs would undermine confidence in science, and negatively affect ratings and audience interest (58). In general, media producers have reported wanting to feature experts who are authoritative, confident, and willing to court controversy--all characteristics which do not necessarily align with effectively conveying scientific knowledge or nuance (59). These are also masculine-coded characteristics and women may receive gendered hostility for displaying them or be more likely to be professionally penalized by senior male colleagues for them, perhaps explaining why women experts are generally more hesitant to appear on television than their male colleagues (59); for more discussion of misogyny in shaping perceptions of female leadership and expertise, see (60, 61).

### Featured experts

Shark Week episodes often repeatedly rely on a subset of hosts/experts; 102 of the 229 hosts/experts were featured in more than one episode. Of those, 80 were featured 2-5 times, 13 were featured 6-10 times and nine were featured more than 10 times. Eight of the nine host/experts were featured in between 10 and 19 episodes, with one person featured in 43 different episodes.

22.7% of the 204 people billed as an expert, scientist, or researcher by Shark Week have no peer-reviewed publications (Fig 6). 14.4% of featured experts have between one and five scientific publications, and although some of these individuals are early career researchers, many are people who are not working professionally in science. For example, one cinematographer is a coauthor on two scientific publications but is primarily not a scientist.

**Fig 6.**
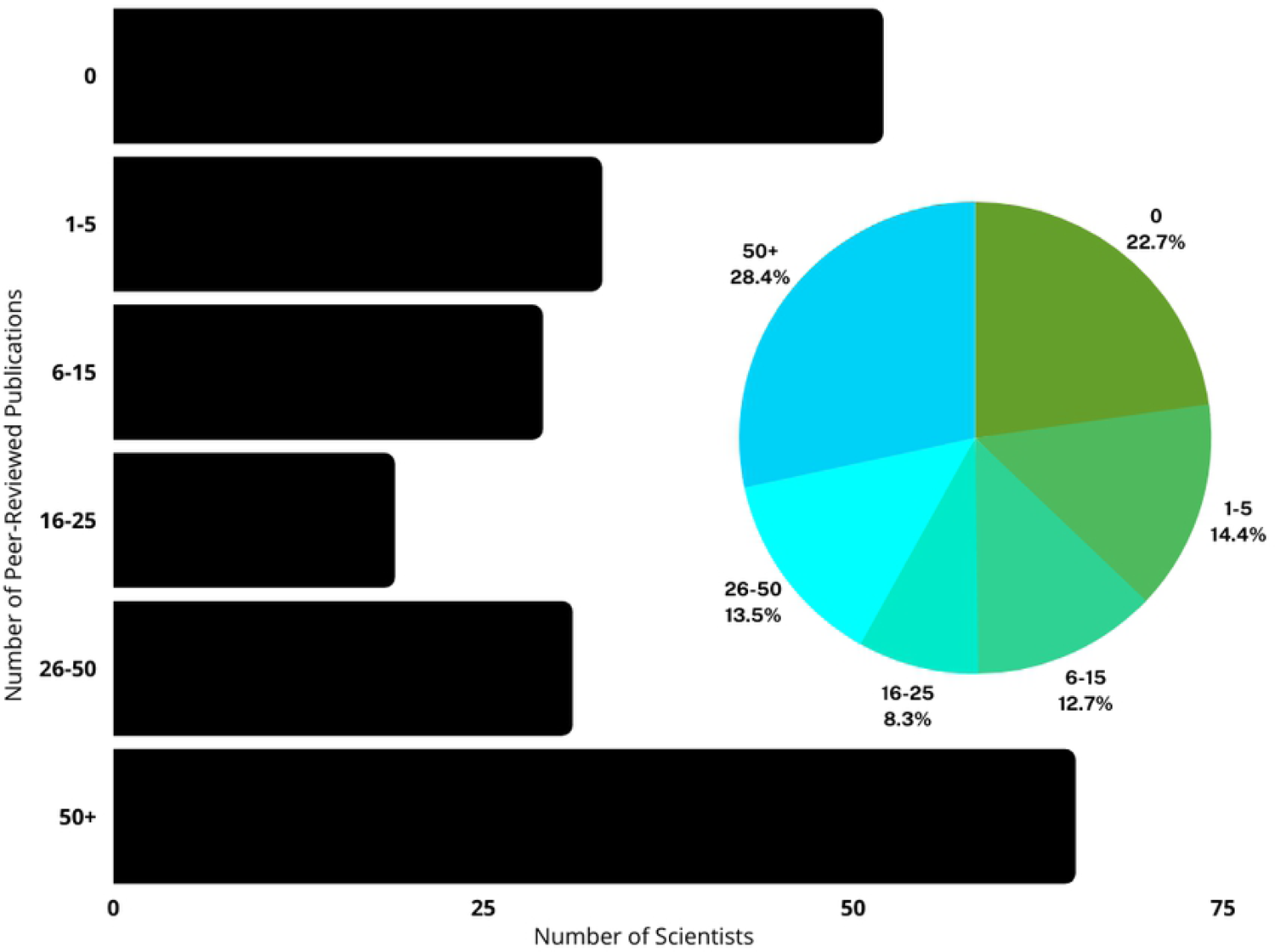
Frequency of number of scientific publications authored by Shark Week experts.

However, just over 41% of experts featured have more than 26 peer-reviewed publications. Although the metric of publications is an imperfect measure of scientific and research contributions, it does provide a general sense of whether someone is actively engaged in publishable scientific research. However, publication metrics and credentials may not be central to television representations of expertise. On U.S. talk shows, experts--particularly “intellectual experts”---are subject to “levelling,” or being treated in ways that present them as equivalently knowledgeable as non-experts. They are often brought on late in an episode, featured alongside non-experts, given little time to speak, frequently interrupted, and may be challenged or disagreed with (62). In some sense Shark Week undertakes a similar leveling process, treating activists, divers, camerapeople and others as having equivalent scientific expertise to credentialed scientists. Of the nine most frequently featured host/experts, three have no peer-reviewed publications, including the host with the most Shark Week episodes (43 episodes).

While there are multiple kinds of useful and relevant knowledge, it may be helpful for Shark Week to more clearly distinguish between scientists and non-scientists (who may well possess other forms of valuable expertise but should not be presented as scientific authorities). It is also noteworthy that many of the most egregious and harmful factual errors or misrepresentations highlighted in criticisms of Shark Week came from non-scientists who Shark Week presents as experts.

Shark Week programming has previously been criticized for overwhelmingly featuring white men as experts in their programming (27) and we were left with the same impression after viewing 201 episodes. 93.9% of experts were perceived by coders as white or white-passing, with only 6.1% of experts perceived as non-white. 24 out of 201 episodes included at least one host/expert perceived by coders as non-white; only one episode included more than one host/expert perceived as non-white. Based on our search, no experts used non-binary pronouns or publicly mentioned being trans*. 78.6% of hosts/experts were associated with male pronouns (an online biography for 2 hosts/experts was not readily available via Google search), whereas the remaining hosts/experts (20.1%) were associated with female pronouns (Fig 7). 60 of the 201 episodes included at least one host/expert associated with female pronouns. Only 11 episodes included more than one host/expert associated with female pronouns; of these nine aired between 2016 and 2018 and one each in 2003 and 2004. Of the 35 experts referred to as “Dr.”, three were associated with female pronouns. We note that two of the male experts referred to as “Dr.” do not have a Ph.D., D.V.M., or similar degree, and that some experts/hosts known to have Ph.D.’s were not referred to as “Dr.” The nine hosts/experts who have been featured in more than 10 episodes are all associated with male pronouns and were perceived as white or white-passing. We note our results over-represent the coverage experts associated with female pronouns and experts perceived by coders as non-white; if a female and/or non-white-passing expert was featured on screen for one episode in an episode where the vast majority of speaking was performed by male and/or white-passing experts, the episode was counted as featuring a female expert.

**Fig 7:**
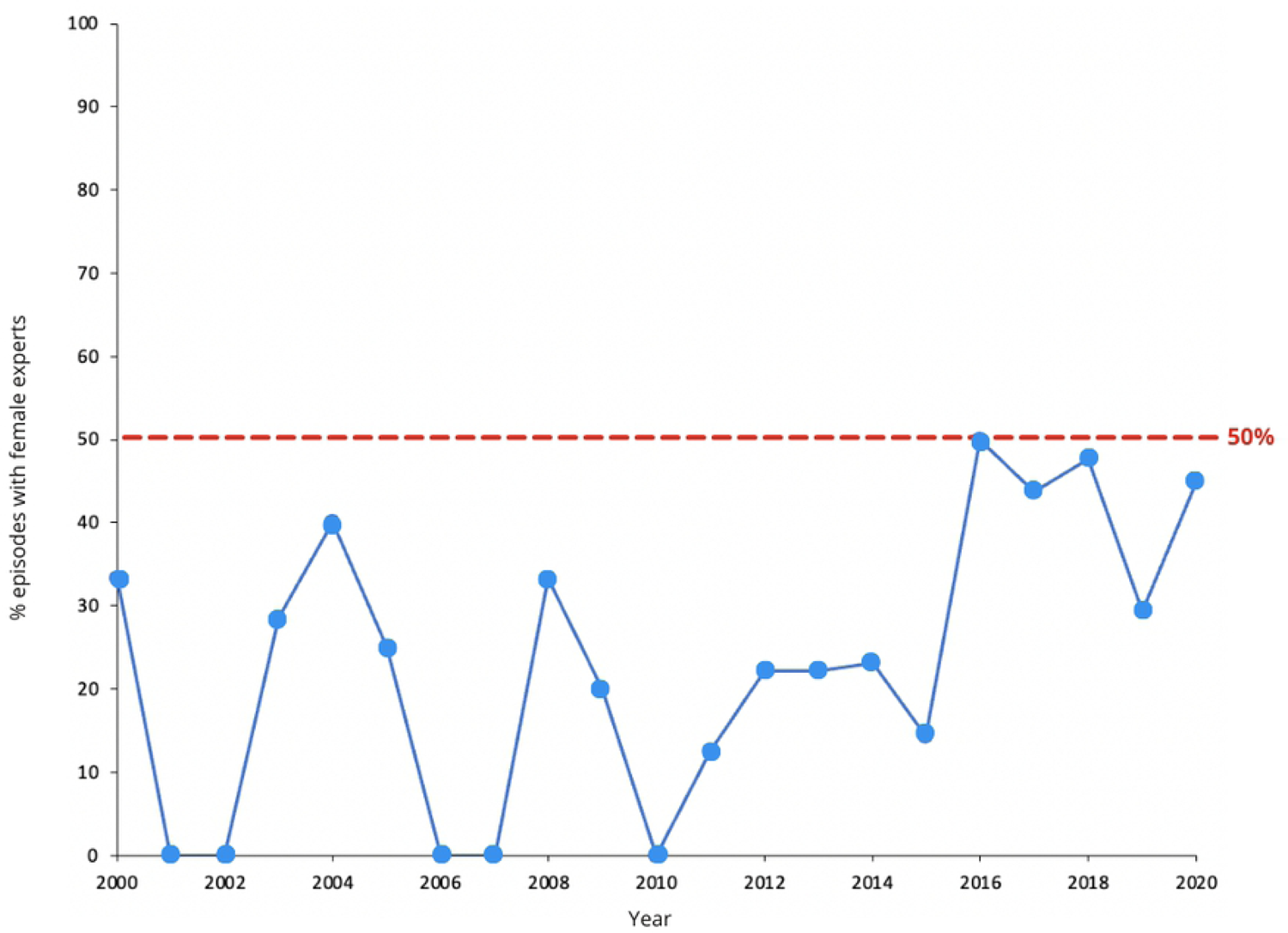
Percent of episodes including any appearance by an expert/host referred to by female pronouns, by year. The red dashed line indicates 50% for a given year. Overall, 20.1% of hosts/experts were associated with female pronouns.

Women are underrepresented in science, filling approximately 26% of jobs, with underrepresentation even more pronounced among women of color (63). STEM (science, technology, engineering, and mathematics) fields in general and shark science in particular are known to suffer from problems with misogyny, harassment, and discrimination (e.g., 56,64,65). Counter-stereotyping and access to same-race and same-sex role models can play an important role in making historically excluded groups feel a greater sense of belonging in science, so availability of role models, including in media, is significant (66–68).

The selection methods for experts appearing on Shark Week have an important influence on content. Experts may be selected for media appearances based on a prior existing relationship with the producer or documentary team, or may be asked to vet or recommend other potential experts being considered (e.g., 69). As people are most likely to have social networks structured around homophily (i.e., primarily composed of people similar to themselves (70)); these recruitment methods can perpetuate a lack of diversity among featured experts. Host/experts are also found through production teams researching social media or published works, such as research papers. This is more likely to favor established, senior researchers with a larger publication record or a higher public profile. It could also result in people with a particularly active social media presence being featured, whether or not they are scientific experts. The limitations created by this recruitment process are not necessarily insurmountable; Shark Week’s chief competitor, National Geographic’s “Shark Fest,” has partnered with the non-profit Minorities in Shark Sciences to improve diversity among their own hosts, while Shark Week has made no such moves publically as of this writing.

### Featured Species

Including species that weren’t the focus of an episode but were briefly introduced by name on screen, at least 79 extant (living) species of shark or species groups (e.g. “hammerhead”, “mako”, “sevengill”, “sixgill”, “thresher”, “wobbegong”) were featured in at least one Shark Week episode (Fig 8, Supplement 3). Additionally, eight extinct species and 13 species of extant non-shark chondrichthyans were also featured (10 batoids (rays), 3 holocephalans (chimera and ratfish)). 46 extant species were featured in more than one episode, 30 appeared in more than five episodes, and 16 appeared in more than ten episodes. Across all episodes, an average of 4.9 species appeared. 39 episodes showed just one species, and 36 of these single-species episodes featured only white sharks (*Carcharodon carcharias)*.

**Fig 8:**
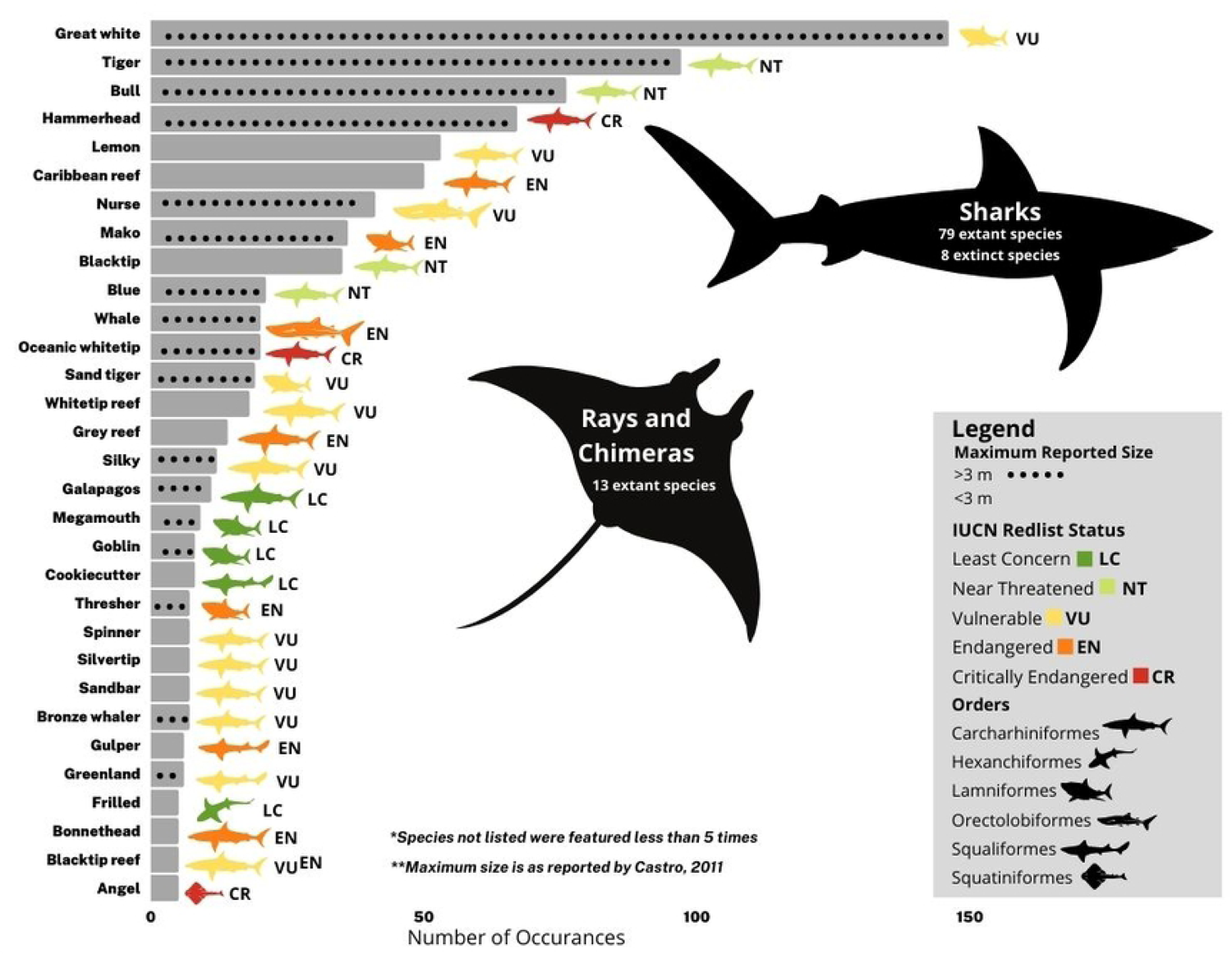
Species appearing in at least 5 Shark Week episodes. Hammerheads were most commonly great hammerheads S. mokarran, though scalloped hammerheads S. lewini and smooth hammerheads S. zygaena were sometimes mentioned. Conservation status reflects that of great and scalloped hammerheads (Critically Endangered); smooth hammerheads are currently assessed as Vulnerable. Mako sharks were almost always shortfin makos Isurus oxyrinchus, but one longfin mako I. paucus was mentioned (both species are Endangered).

Across the 201 coded episodes, the most common species featured were white sharks *C. carcharias* (18.4% of all episodes), tiger sharks *Galeocerdo cuvier* (12.2% of all episodes), bull sharks *Carcharhinus leucas* (9.6% of all episodes), and hammerhead sharks Sphyrnidae (8.4% of all episodes) (Fig 8). Often the specific species of hammerhead was not mentioned, so all hammerheads were grouped together for analysis; when species were specified, great hammerheads *Sphyrna mokarran* were featured most often (62.3%) with the occasional scalloped (21.7%) and smooth (2.9%) hammerheads (*S. lewini* and *S.zygaena*, respectively).

The species highlighted show some interesting contrasts with similar analyses of shark species of interest in scientific publications (57) and popular press (19) coverage. While white sharks appear in the top five featured species in Shark Week, scientific publications, and popular press coverage, some of the most-studied species (bonnethead shark *Sphyrna tiburo*, sandbar shark *Carcharhinus plumbeus*, and spiny dogfish *Squalus acanthias*) are rarely featured in any Shark Week episodes (5, 7, and 1 episode, respectively) (Supplement 3). Similarly, some of the species that received the most media attention in popular press articles (19) such as the porbeagle *Lamna nasus* and basking shark *Cetorhinus maximus* were rarely featured in any Shark Week episodes (2 and 4 episodes, respectively)(Supplement 3). As in other forms of popular media, more highly threatened species were not more likely to be featured, with an overall tendency to large, charismatic, well-known species.

### Featured localities

Though dozens of countries were featured in at least one episode each, a handful of filming locations dominated. The United States was the most common filming location (24.2% of all episodes), followed by the Bahamas and South Africa with 15% each, and New Zealand, Australia, and Mexico with approximately 10% each (Fig 9). Within the United States, 31.5% of episodes took place in California, followed by Florida (26.7% of episodes), Hawaii (17.8% of episodes), and Massachusetts (9.6% of episodes). At least one episode took place in nearly every coastal state’s waters (except for Delaware, and noting that most shows featured in Georgia were filmed at the Georgia Aquarium). However, most states other than Florida, California, Hawaii, and Massachusetts were not featured often, and some states were featured only once.

**Fig 9.**
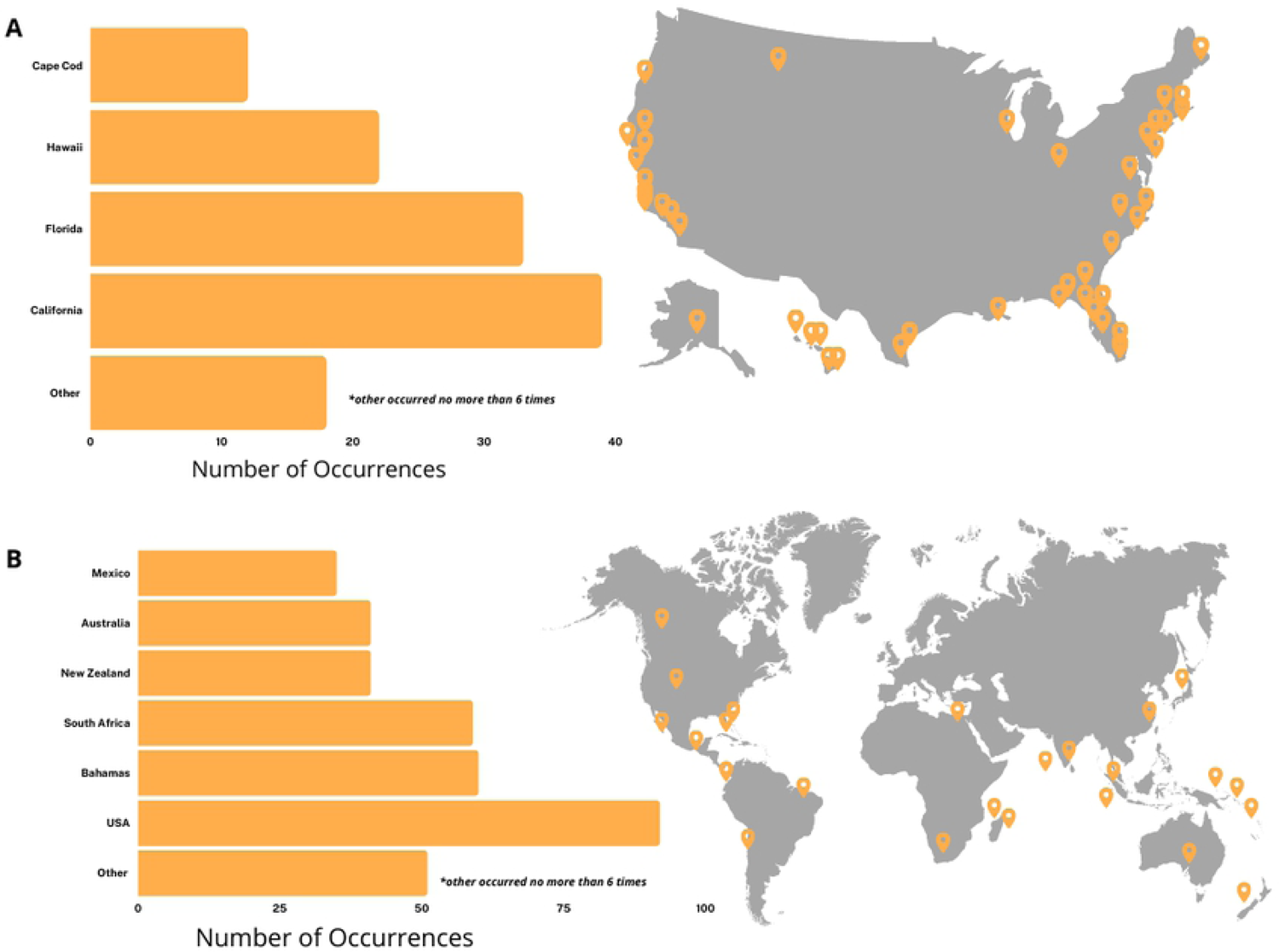
Shark Week filming locations. (A) Locations by country; (B) Locations within the United States.

This geographic focus on just a few countries (and on a relatively small number of locations within those countries) in part reflects a focus on particular species and researchers, though Shark Week episodes regularly feature experts who have no particular experience with a location, but fly in to the location just for filming the episode. Although sharks are circumglobal, familiar and logistically simple sites in which filmmakers have prior experience or existing relationships may be favored for filming (71), potentially acting as a factor which contributes to reducing the diversity of species, locations, narratives, and scientists featured. For example, two of the top three filming localities have majority Black populations; 91% of the Bahamian population is Black (72) and approximately 80% of the South African population is Black African (73). Despite 30% of filming localities being located in these two countries, non-white experts are rarely featured in Shark Week (this study; 27).

#### Messaging

174 (86.6%) of the coded episodes had a stated goal at the beginning of the episode. Of these episodes, 64 (36.8%) did not address their stated goals during the course of the episode, and 110 (63.2%) claimed to have accomplished their stated goal. The stated goals varied from specific research goals to answering general questions about shark behavior (see Supplement 2 for examples). If the goal of these episodes is to educate viewers, it is important that they have a clearly stated purpose and that this purpose is addressed. The fact that this often did not happen shows that many episodes serve no purpose beyond imagery of sharks.

148 (73.6%) of the coded episodes included some sort of fear-mongering language or negative portrayal of sharks. These comments mostly focused on shark bites on humans (Table 1). On the other hand, 127 (63.2%) of the episodes had at least one mention of sharks as awe-inspiring, beautiful, misunderstood, or ecologically important (Table 2). Notably, this was often a brief mention that played over the ending credits, while fear-mongering-type narrative often occurred throughout the episode.

**Table 1:**
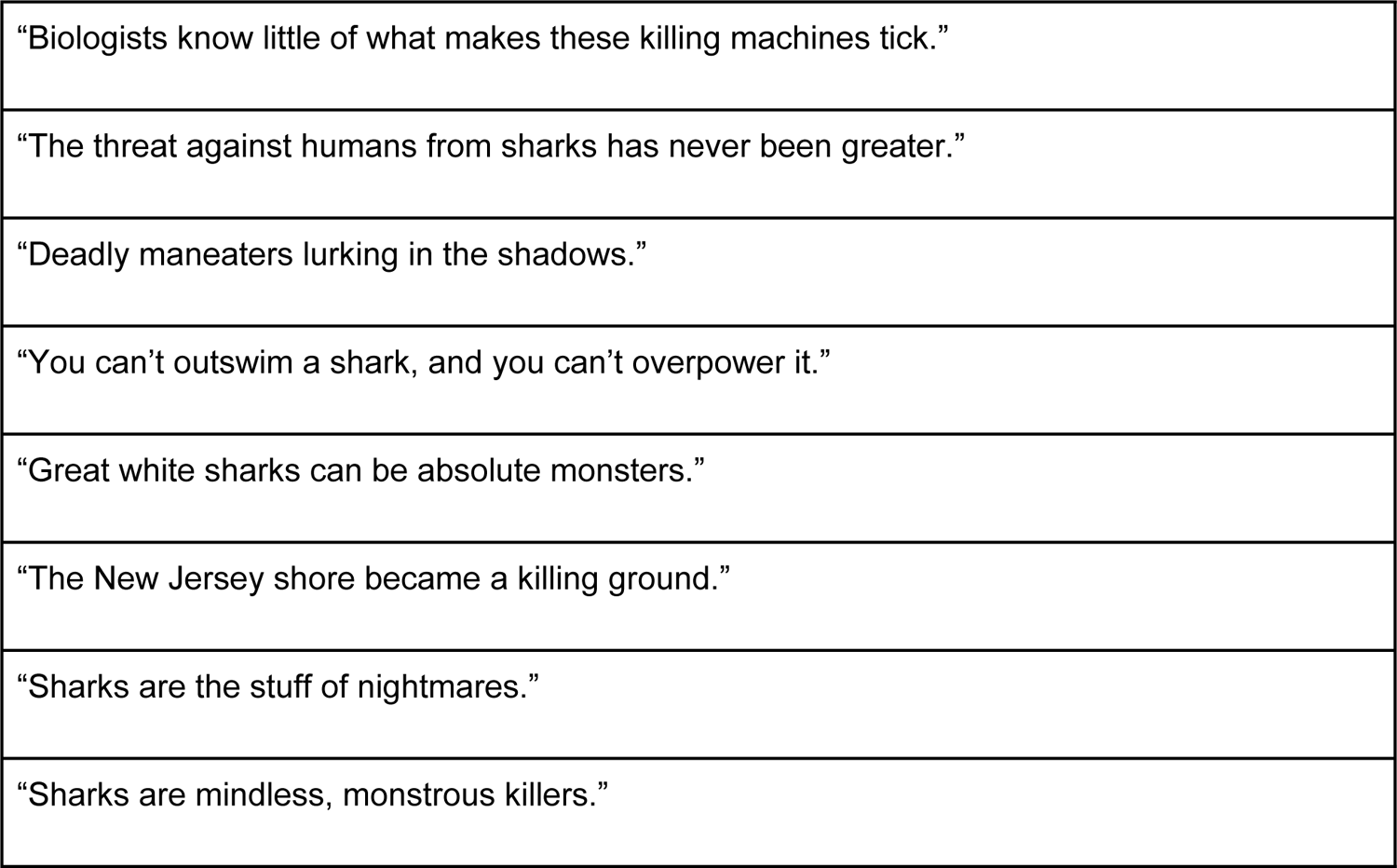
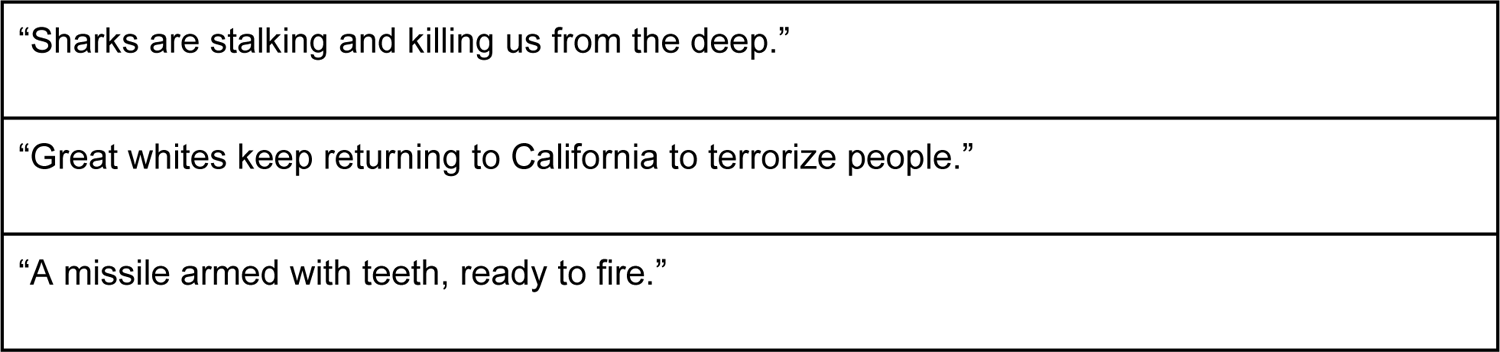
Representative example dialogue and narration showing sharks in a negative light.

|  |
| --- |
| "Biologists know little of what makes these killing machines tick." |
| "The threat against humans from sharks has never been greater." |
| "Deadly maneaters lurking in the shadows." |
| "You can't outswim a shark, and you can't overpower it." |
| "Great white sharks can be absolute monsters." |
| "The New Jersey shore became a killing ground." |
| "Sharks are the stuff of nightmares." |
| "Sharks are mindless, monstrous killers." |
| "Sharks are stalking and killing us from the deep." |
| "Great whites keep returning to California to terrorize people." |
| "A missile armed with teeth, ready to fire." |

**Table 2:** Representative example dialogue and narration showing sharks in a positive light.

|  |
| --- |
| "The balance and health of the ocean depends on their survival." |
| "A marine ecosystem is not healthy without top predators." |
| "You cannot remove the top predators without affecting every link below." |
| "Sharks are misunderstood animals." |
| "Sharks are more valuable alive than dead." |
| "Gentle giants" |
| "Great white sharks are one of the most awe-inspiring animals on the planet." |
| "Sharks are clearly intelligent, not mindless killers." |
| "Seeing a great white breach is one of the most spectacular things in nature." |
| "What's not to love?" |

The language used to describe sharks does matter, as studies have shown that negatively valenced words like “attack” can contribute to negative public sentiment towards sharks (49, 74). Public acceptance of predators is related to the frequency and intensity of interactions (especially negative interactions) with humans, so support for shark conservation is likely to be related to perceived frequency of bites or human injuries (i.e., “attacks”) (75). Media portrayal of these issues has indeed been shown to play a role in public support for shark conservation (11). Muter et al. (14) and Neff and Hueter (74) also found that news stories about sharks largely focus on fear-mongering and exaggerated stories of sharks biting people rather than on shark research or conservation. Exposure to violent Shark Week programming has been shown to induce greater levels of fear of sharks (76), and fear correlates with support for policies, like shark culls or beach netting, that are harmful to shark conservation (77).

One area in which Shark Week programming may be effective at reducing fear is through episodes including neutral and positive interactions with sharks, which have been shown to improve public perceptions (78). Episodes of Shark Week in the last several years typically include at least one well-known celebrity interacting with sharks (e.g., Shaquille O’Neal, Will Smith, Adam Devine, Ronda Rousey, Craig Ferguson). Celebrity actions, opinions, and endorsements are known to influence the attitudes we adopt and the decisions we make (79), and in conservation specifically, celebrity endorsement of a cause yields higher willingness-to-engage amongst the public (80). However, it should be noted that Shark Week’s celebrity episodes often feature unnecessary artificial danger or inappropriate interactions with animals such as chasing, riding, or harassing them, which could undermine any positive messaging and potentially endanger people’s safety (81).

In terms of specific threats to sharks and shark conservation, 28 episodes (13.9%) mentioned shark finning or the shark fin trade, and eight (4.0%) mentioned that people eat shark meat.

However, Shark Week episodes are generally lacking in actionable educational content about shark conservation. 107 episodes (53.2%) at least briefly mention something related to conservation, often vague statements about shark population decline, the ecological importance of sharks, or extinction risks. Just six episodes (3.0%) mentioned anything specific about shark conservation policy or specific ways that Discovery’s audience could help; these statements were mostly about individuals choosing to not eat shark fin soup or releasing sharks they catch. There was no content encouraging viewers to speak to government officials about specific ongoing policy discussions, advising them to avoid specific seafoods with shark bycatch, requesting donations to nonprofits that have a track record of success, or incorporating any other common advice given by experts to those who want to help conserve sharks. When suggestions are provided during programming, many of them are not actionable in any way that could actually be useful to shark conservation efforts, an enormous missed opportunity given Shark Week’s massive audience and the general lack of public pro-sustainability engagement in US shark fisheries discussions (82). Past attempts to leverage their audience included a 2014 social media ad with five ways that people could help sharks, which included “report shark attacks” and “avoid shark fishing in marinas” as suggestions without explanation. The most specific was “lobby for shark protection,” but no information was provided on who to lobby or what to ask them to do. Exposing a large audience to vague platitudes is of questionable value for conservation and may even undermine existing campaigns (82).

### Expert attitudes toward Shark Week

A survey of expert shark scientists revealed broad concerns about the role of media misinformation in general, and Shark Week specifically, in perpetuating misinformation about shark research and conservation. 102 experts responded to the survey, but not everyone responded to every question, and therefore the following percentages are relative to the number of respondents that answered the particular question. Survey respondents generally believe that public understanding of sharks is a significant factor influencing their conservation. 64% (N=49) of respondents refer to public attitudes towards sharks as significant/important or very significant/very important to shark conservation. Survey respondents reported being very concerned about how sharks are portrayed in the media, a primary way that the public becomes informed about environmental issues. 86% of responses to this question (N=74) reported believing that mainstream media coverage of shark related issues is not factually accurate, and while the question did not specifically ask about Shark Week, many respondents brought up their concerns with sensationalist and inaccurate coverage included on Shark Week unprompted (Table 3).

**Table 3:**
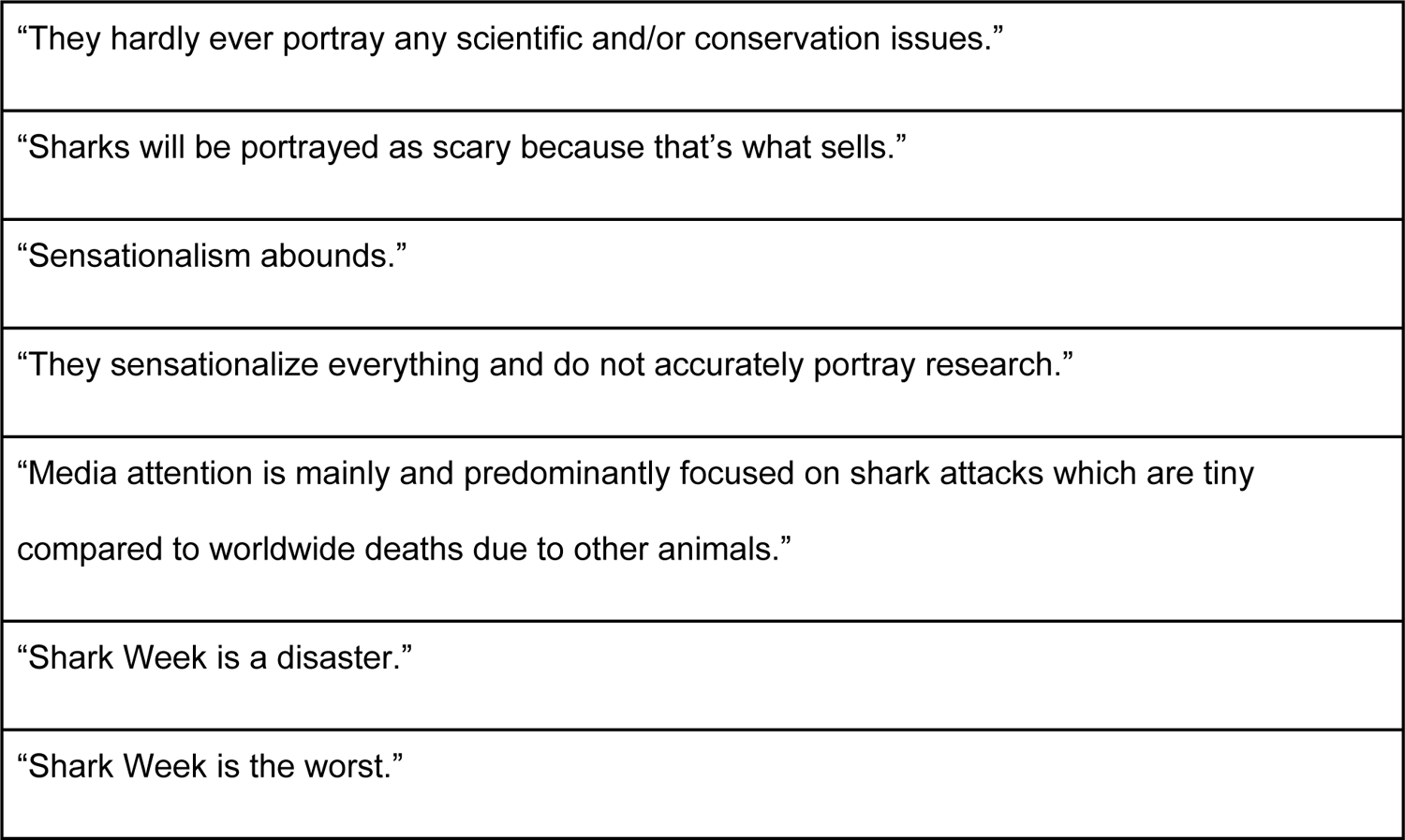
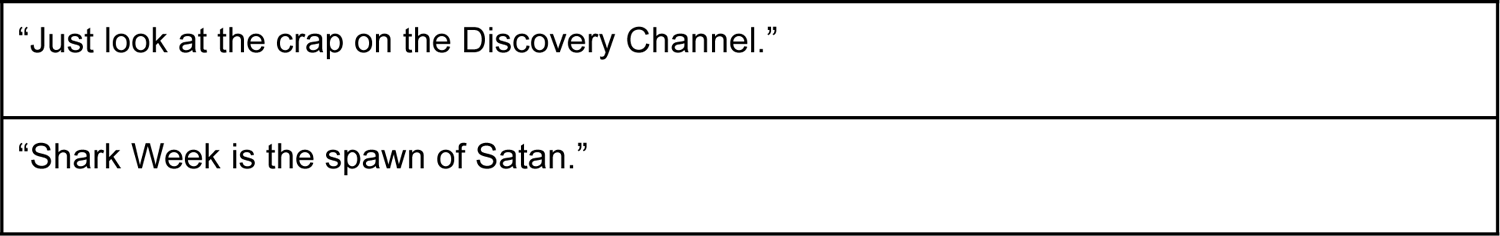
Selected responses to the survey question “In your opinion, does the mainstream media accurately portray shark science and conservation issues” demonstrating survey respondents’ concerns about how shark science and conservation issues are portrayed in the media in general, and about Shark Week specifically.

The disconnect between scientists and the producers of science-related programming may help explain the lack of support for Shark Week among a significant portion of professional shark scientists. In general, scientists report that much of what is called science on television is not science but entertainment, while producers tended to define science programming much more broadly and believe television represents science well (58). Scientists who do appear on television emphasized the particular additional skill sets required, while the broader consensus from the scientific community seems to be that scientists are “…better off doing science, but letting the broadcasters do the science stories’’ (58, p. 129). This, however, presents obvious challenges for programming such as Shark Week, where scientific inaccuracy is already a major source of complaint and conflict--many scientists both want to see programming improved and do not want to have to be involved in those improvements. This caution is understandable, given recent fictional programming and the reality that some scientists have reported being misled and misrepresented by Shark Week producers (29). However, it is unlikely that programming will improve without the continued participation and engagement of scientists.

### Recommendations

The scale of Shark Week’s platform to communicate with the public about sharks means that even minor adjustments to programming could have a meaningful effect. Some of the simplest improvements involve reducing harmful sensationalism (including perceptions of the dangerousness of many activities), enhancing factual accuracy and raising editorial standards, and clearly distinguishing between fact-based and fictional programming. Similarly, explicit differentiation between credentialed scientific experts and non-scientist hosts would be helpful in avoiding inadvertently legitimizing incorrect information.

In portrayals of science and scientists, it would be helpful to feature real science and more realistic scientific methods (even if recreated or dramatized), a wider range of shark species, and a more diverse range of scientists. These changes would likely help with factual accuracy while also benefiting the diversification of shark science, recruitment in STEM, and public recognition of the work of scientists from historically excluded groups.

In terms of the effects these changes might have on the public, some studies of students of varying ages have shown that increasing knowledge about animals increases positive attitudes towards those animals (83, 84), including for sharks in particular (85). Television has the potential to drive conservation action or intention--for example, an increase in internet searches for conservation charities and sustainable practices were seen during and after the airing of *Blue Planet II* episodes (86, 87). This does not mean that providing the public with positive representations of sharks, or accurate information about them, represents a solution to conservation problems or will necessarily generate interest or concern about them in itself (88–90). However, evidence suggests playing on existing negative stereotypes--even with an intent to challenge them--can actually serve to reinforce them (e.g., 91). Best practices for improving the public image of sharks include shifting away from negative stereotypes and providing detailed information about how conservation problems connect to people’s lives and what actions they can take to help (78).

The majority of Shark Week episodes do contain at least some educational content about sharks (often vague or brief mentions), with most episodes falling into the (broad) categories of Research or Natural History (Fig 3). Even episodes focused on bites or attacks can offer some educational value when they include scientifically accurate information, though this is often undermined by conflicting messages and sensationalism, as in programs which terrorize viewers and then briefly mention shark conservation as the credits roll. While conservation content may not be appropriate for every episode, providing actionable steps for viewers is necessary in order for them to move from positive attitudes towards behavior that supports shark conservation; viewers who are misinformed or under-informed about key issues related to conservation are unlikely to support expert-backed policy solutions in useful ways (19). For example, a small number of episodes correctly connected shark fishing for meat and fins to the current population decline of sharks (3, 7). However, no episodes linked these facts to specific actions the audience could take to make a difference, and only six episodes included anything arguably specific and detailed about conservation.

## Conclusion

Shark Week has a complicated history over the course of its 30+ years, and has received substantial criticism for scientific inaccuracy, while also unquestionably increasing the public attention paid to sharks. This analysis attempted to quantitatively assess some of the trends and practices seen in Shark Week programming that have been anecdotally discussed for decades.

Our analyses demonstrate that the majority of episodes are not focused on shark bites, although such episodes are common and many titles and episodes are framed around fear, risk, and adrenaline. Including tangential mentions, a surprising number and diversity of shark species have been featured, although anecdotal descriptions of disproportionate attention to particular large charismatic species are supported by our data. Shark Week’s depictions of research and of scientists are biased towards particular research methodologies and (mostly white, mostly male) scientists, including non-scientists being presented as scientific experts even as they share incorrect information. Results suggest that as a whole, Shark Week is likely contributing to collective perceptions of sharks as monsters, and that even relatively small alterations to programming decisions could substantially improve the presentation of sharks and shark science and conservation issues.

This requires a complex balance of Shark Week’s potentially competing goals to educate and entertain audiences and contribute to conservation. If Shark Week does not retain viewers, any efforts to improve programs’ educational and conservation impact will not be meaningful. These competing imperatives have been recognized since at least the 1940s with radio shows such as *Great Moments in Science* and television’s *The Nature of Things* (LaFollette 2008) successfully combining entertainment and accurate educational content. Successful, scientifically accurate programming featuring Don Herbert (“Mr. Wizard”), Carl Sagan, and Bill Nye, among others, succeeded because hosts displayed excellent and entertaining communication skills (92, 93).

Programming featuring stunning visuals and music such as *Blue Planet* and *Planet Earth* effectively entices viewers with ‘visual and aural pleasure’ (*sensu* 94) while also delivering accurate educational information.

Given its popularity and global viewership, Shark Week has the potential to generate interest in both sharks and scientific careers among viewers. However, Shark Week fails to feature the full range of shark research topics and methods and the diversity of people performing research on sharks. Cultivating a positive attitude toward sharks through Shark Week has the potential to drive enhanced support of shark and ocean conservation efforts. Currently, through a series of unnecessary and harmful programming choices, Shark Week can be seen as a missed opportunity to benefit sharks, shark science, and shark conservation.

## Acknowledgements

This research was supported by funding from Allegheny College. We thank J. Carrier and R. Hueter for providing personal copies of episodes for our use. We also thank R. Kajiura for providing his professional independent analysis of episode titles and A. Knupsky for pointing us toward the ANEW. C. Bailey and B. Davis gave invaluable advice on demographic analysis. This study was inspired by conversations held in Allegheny College’s 2019 FS101 course “Sharks and Recreation”, taught by author LBW - thank you to the Sharks and Rec students for their conversations and insights.

## Supporting information

S1. Shark Week episodes by year. Blue column indicates whether the episode was included in the content analysis. Yellow columns indicate titles deemed negative using the ANEW, using context, and with the ANEW and context combined.

S2. Coding guidelines for the content analysis of Shark Week episodes.

S3. Chondrichthyan species featured in Shark Week episodes and number of episodes.

